# A robust closed-tube method for resolving *tp53*^M214K^ genotypes

**DOI:** 10.1101/2023.01.20.524903

**Authors:** Katherine M. Silvius, Genevieve C. Kendall

## Abstract

The *tp53*^M214K^ zebrafish mutant developed by Berghmans et al 2005^1^ is a versatile platform with which to model a diverse spectrum of human diseases. However, currently available genotyping methods for this mutant require lengthy processes such as restriction digests and outsourced Sanger sequencing. To address this deficiency, we leveraged high-resolution melting analysis (HRMA) technology in conjunction with a parallel, in-tandem wildtype spike-in approach to develop a robust genotyping protocol capable of discriminating *tp53*^M214K^ zygosity. Here, we describe our method in detail. We anticipate that our genotyping protocol will benefit researchers utilizing the *tp53*^M214K^ zebrafish mutant by offering reliable results with a faster turnaround time than conventional approaches.

## Main text

Animal models with *TP53* mutations are critical for the development of *in vivo* genetic systems that recapitulate the innate complexity of the human disease spectrum. *TP53* mutations are often required for zebrafish models to form tumors, as demonstrated with the BRAFV600E driven melanoma and PAX3/7-FOXO1 fusion driven rhabdomyosarcoma zebrafish models.^2,3^ To these ends, the *tp53*^M214K^ zebrafish mutant developed by Berghmans et al 2005^1^ (also referred to as *tp53*^zdf1^) has been widely used when building vertebrate disease models that require a *TP53*-mutated background. This mutant is commercially available (ZIRC, ZL1057) and harbors a missense mutation in the DNA-binding domain of *tp53*, which mimics the most common type of *TP53* mutation found in humans.^1,4^ Specifically, the mutant allele contains a T>A point mutation that results in a change from methionine to lysine and consequential dysregulation of *tp53* function. Spontaneous tumors begin to form around 8.5 months in homozygous fish.^1^ Since zebrafish are not in-bred, the workflow of line maintenance necessitates efficient genotyping to support research efforts. Currently, stereotyped approaches for genotyping the *tp53*^M214K^ mutant are limited to multi-step PCR-based assays that often require PCR clean-ups and resolution by sequencing or gel, which hinders the genotyping scale and turnaround time (Fig. 1A-B).^1,5,6^

**Figure 1:**
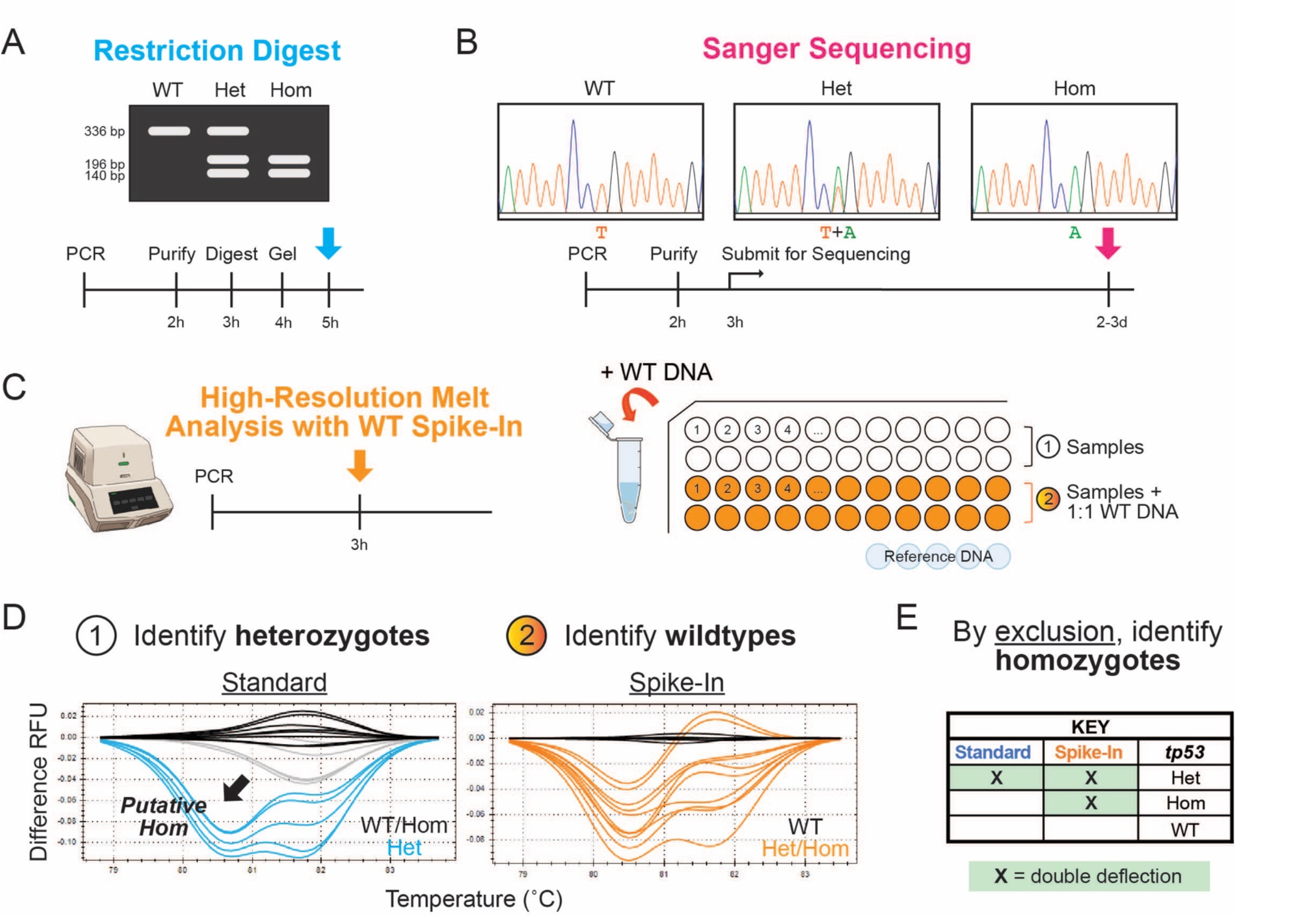
An improved workflow for *tp53*^M214K^ genotyping. Conventional genotyping protocols for the *tp53*^M214K^ mutant like **(A)** restriction digest and **(B)** Sanger Sequencing based approaches are time-consuming and require multiple steps, whereas our protocol predicated upon **(C)** high-resolution melt analysis with a wildtype spike-in requires a single PCR program on a standard qPCR machine (*left panel*), with samples that are run in parallel with and without a 1:1 wildtype spike-in alongside a set of wildtype reference DNA for comparison (*right panel*). **(D)** The analysis strategy utilizes the characteristic double deflection pattern of *tp53* heterozygotes to first identify heterozygotes in the standard run without the spike-in. Here, only about 60% of homozygotes can be called. Homozygotes verified by sequencing are labeled in gray, and those that can be called by only the standard HRMA are indicated by a black arrow (*1*). Since wildtype DNA added to homozygotes yields a similar pattern to allelic heterozygotes, samples that do not demonstrate that deflection pattern in samples with the spike-in can be identified as wildtypes (*2*). **(E)** True homozygotes can then be identified by comparative exclusion as illustrated in the table. Abbreviations: WT, wildtype; Het, heterozygote; Hom, homozygote.

Here, we sought to leverage high-resolution melt analysis (HRMA) technology to decrease the time and steps required for genotyping *tp53*^M214K^ mutants (Fig. 1C). HRMA is a technique that incorporates a gradual, systematic heating step immediately following amplification. Zygosity is then discriminated based upon differences in DNA melting temperature of the mutant alleles compared to a wildtype reference. HRMA offers many advantages over conventional genotyping approaches, such as minimal set-up, a single closed-tube reaction, and simple analysis.

Additionally, HRMA is sensitive enough to detect single-nucleotide polymorphisms (SNPs) and has been successfully utilized in zebrafish, including genotyping a different point-mutation *tp53* mutant.^7^ However, a detailed protocol has not been established for the frequently used *tp53*^M214K^ line (see Supplemental Material for detailed protocol).

To begin, we designed^8^ and tested a series of primer sets flanking the *tp53*^M214K^ mutation site with amplicon sizes ranging from 63-125 bp. Amplicons of <100 bp are often used for SNP scanning as small changes in the mutant allele are amplified when the total region is also small.^9^ However, we were not able to consistently differentiate *tp53*^M214K^ genotypes.

Heterozygotes were always clearly defined by a characteristic double deflection pattern, but repeated tests demonstrated that a small but persistent percentage of homozygotes often clustered with wildtypes (Fig. 1D). This is likely due in part to the fact that T>A is associated with the smallest energy change of all possible point mutations, which inherently makes *tp53*^M214K^ zygosity more difficult to discriminate.^9^ Additionally, as the primers move closer to the mutation site, the GC percentage continually increases (>60%), which precludes the feasibility of continuing to decrease the amplicon size.

To address this challenge, we took the four most promising primer sets (110-bp, 96-bp, 83-bp, and 70-bp) and titrated the DNA input by serial dilution from 300 ng to 12.5 ng. While this showed that the assay, particularly with the 96-bp primer, is able to detect all heterozygotes and most homozygotes in a working range of 10-200 ng DNA input, we were still unable to achieve the consistency necessary for a robust, stereotyped genotyping protocol.

We then decided to test our 96-bp HRMA assay with a complementary wildtype spike-in, which is similar to an approach previously used to genotype this mutant, although that method was not reported in detail.^10^ Evaluating samples in the same HRMA run both with and without wildtype DNA spiked in should allow for genotype resolution by comparative exclusion. For example, adding wildtype DNA will drop a homozygote’s deflection to that of an easily identifiable heterozygote without affecting wildtype samples, and true heterozygotes can be pulled out from the run without the spike-in (Fig. 1D-E). We confirmed this by running samples at 150 ng total DNA with and without a 1:1 wildtype spike-in. In blinded tests without the spike-in, only ∼60% of homozygotes could be accurately called, but when the parallel spike-in was added, all genotypes were called correctly (Supplemental Material).

Additionally, since our assay showed promise at a wide range of DNA concentrations in initial tests, we decided to test the requirement for normalization. Using a standard 1 µL DNA volume rather than a standard ng DNA input greatly reduces set-up time. Although the concentration of a typical DNA sample derived from a tail-clip varies, we were able to achieve comparable results using samples ranging from 100 to 400 ng/µL with the same high level of confidence without needing to first normalize the DNA, thereby increasing the throughput of our assay Altogether, we present an HRMA-based genotyping protocol which leverages a complementary, in-tandem wildtype spike-in approach to accurately resolve *tp53*^M214K^ zygosity. We describe this protocol in detail in the supplemental material as a faster alternative to conventional digest and sequencing based approaches.

## Acknowledgments

We thank Dr. Raphael Malbrue, Dr. Laurie Goodchild, Logan Fehrenbach, and the other Animal Resources Core Zebrafish Facility team members for exceptional care and collaboration in maintaining our zebrafish colony. We thank Dr. Matthew Cannon, Dr. Matthew Kent, Jack Kucinski, Delia Calderon, Emma Harrison, and Amanda Jay for their technical support and helpful discussion.

## Author Contributions

G.C.K. and K.M.S. conceptualized and designed the project. K.M.S. conducted the experiments, analyzed the data, and wrote the original draft. G.C.K. supervised the project, acquired the funding, and reviewed and edited the manuscript.

## Animal Ethics Statement

Zebrafish are housed in an AAALAC-accredited, USDA-registered, OLAW-assured facility in compliance with the Guide for the Institutional Care and Use of Laboratory Animals. All research procedures involving animals are approved by the IACUC at The Abigail Wexner Research Institute at Nationwide Children’s Hospital (AR19-00172).

## Supplemental Materials

### Genomic DNA Isolation

1. Anesthetize juvenile or adult zebrafish in 0.2 mg/mL Tricaine diluted in fish water and use a sterile razor to cut the tail fin just before the cleft where the tail bifurcates into the two lobes.
2. Transfer the tail-clips to a PCR strip tube or a 96-well plate.
3. Add 100 µL of Lysis Buffer (10mM Tris-HCl pH 8.3, 50mM KCl, 0.3% Triton X-100, 0.3% NP40) to each tail-clip.
  a. For 50 mL Lysis Buffer: Add 500 µL 1M Tris-HCl pH 8.3, 2.5 mL 1M KCl, 150 µL Triton X-100, and 150 µL NP40 to 47 mL RO water. Shake and/or gently heat in a water bath to dissolve detergents. Store at 4°C.
  b. Transfer samples to a thermocycler and run program with a 95°C lid: 98°C x 10 min, 12°C x 10 min, END.
4. Add 10 µL Proteinase K, 10 mg/mL, to each fin, flick or vortex to mix, then spin down.
5. Return samples to the thermocycler and run program with a 95°C lid: 55°C x 1.5-16 hours, 98°C x 10 min, and HOLD at 12°C.
6. Store DNA at 4°C overnight or proceed directly to the HRMA protocol.

### Wildtype References

High-resolution melt analysis (HRMA) requires samples to be compared to a set of known wildtype references in order to set the baseline melting profile for mutant allele detection. In these studies, AB fish were used as the reference since AB is the wildtype background of the *tp53*^M214K^ line. At least 5 known wildtype samples were run per plate so that up to 2 samples with an unexpected or abnormal curve in the reference cohort could be excluded without adversely affecting downstream analysis. Wildtype reference DNA can be collected in bulk using the standard practices described above and kept at -20°C for longer-term storage.

### Instrumentation and Software Requirements

- Real-time PCR machine: The HRMA software is compatible with all Bio-Rad CFX real-time PCR systems. Here, we used a CFX384. Each machine that will be used for HRMA must first be calibrated to ensure that the dye in the Precision Melt Supermix (Bio-Rad, #1725112) is correctly detected. This is done with the Melt Calibration Kit (Bio-Rad, #1845020) and only needs to be performed once per machine.
- CFX Maestro Software for Windows PC (Bio-Rad, #12013758): This program is only used to convert the run data from the PCR machine to a file type that can be processed using the HRMA software.
- Precision Melt Analysis Software (Bio-Rad, #1845015), 2 user licenses: There is an available option that packages one calibration kit with the purchase of the software.

### Note on Genotyping Call Efficiency

When only the standard run with 1 µL DNA is evaluated, all heterozygotes are called with confidence. If only heterozygote mutant scanning is desired, the standard set alone is sufficient. However, a similar strategy resolves only ∼60% of homozygotes (i.e., samples that deflect lower than -0.04). These are true homozygotes but the total number is often underestimated whereas the wildtype population is overestimated, denoted in pink below. However, when the complementary corresponding spike-in samples are evaluated as well, all calls are correct.

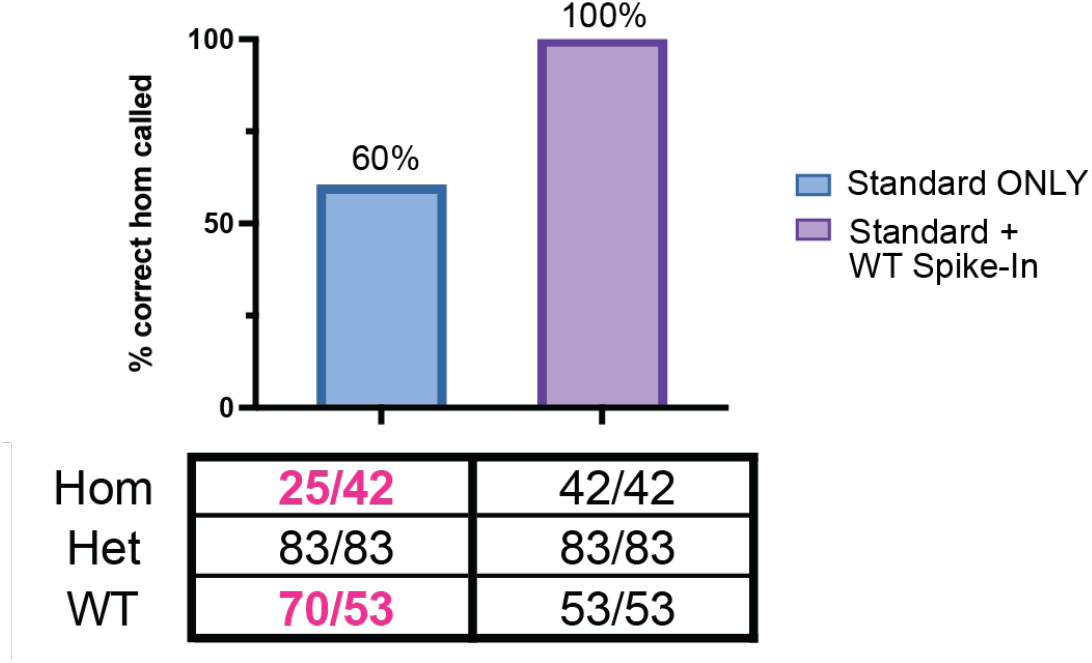

### Detailed *tp53*^M214K^ HRMA Genotyping Assay

#### Materials

- Precision Melt Supermix (Bio-Rad, #1725112)
- tp53 HRMA FWD Primer: GGACAACTGTGCTACTAAACTACATG
- tp53 HRMA REV Primer: CCTGAGTCTCCAGAGTGATGATT
- 384-well plate (Bio-Rad, #HSP3805)
- Microseal ‘B’ PCR Plate Sealing Film (Bio-Rad, #MSB1001)

#### Protocol

1. Prepare a master mix scaled appropriately to account for the total number of samples: **total** = (# unknown) x 2 + (4-5 wildtype references) + 1 negative control + extras

2. Add 19 µL master mix per well of a 384-well plate (note: 96-well plate would work too).
3. Briefly spin down all DNA samples in a mini-centrifuge.
4. For the wildtype references and the standard set of samples without the spike-in, add 1 µL DNA. For the corresponding set with the complementary spike-in, first add 0.5 µL sample DNA, then go back and add 0.5 µL wildtype DNA to each well. *Make sure to avoid any debris at the bottom of the genomic DNA preps*.
5. Cover the plate with B adhesive film and roll to completely seal. *Avoid touching the top*.
6. Briefly vortex and microfuge in a plate spinner.
7. Run the following HRMA program on a real-time PCR machine with 95°C lid and 20 µL reaction volume: 95°C x 3 min, (95°C x 15 sec, 60°C x 20 sec, Plate Read, 70°C x 20 sec) x 44, 65°C x 30 sec, melt curve 65°C-95°C with 0.2°C/step for 5 sec plus Plate Read, 95°C x 15 sec.

| COMPONENT | 1X |
| --- | --- |
| Precision Melt Supermix | 9 µL |
| *tp53* HRMA FWD, 10µM | 0.5 µL |
| *tp53* HRMA REV, 10µM | 0.5 µL |
| Nuclease-free water | 9 µL |

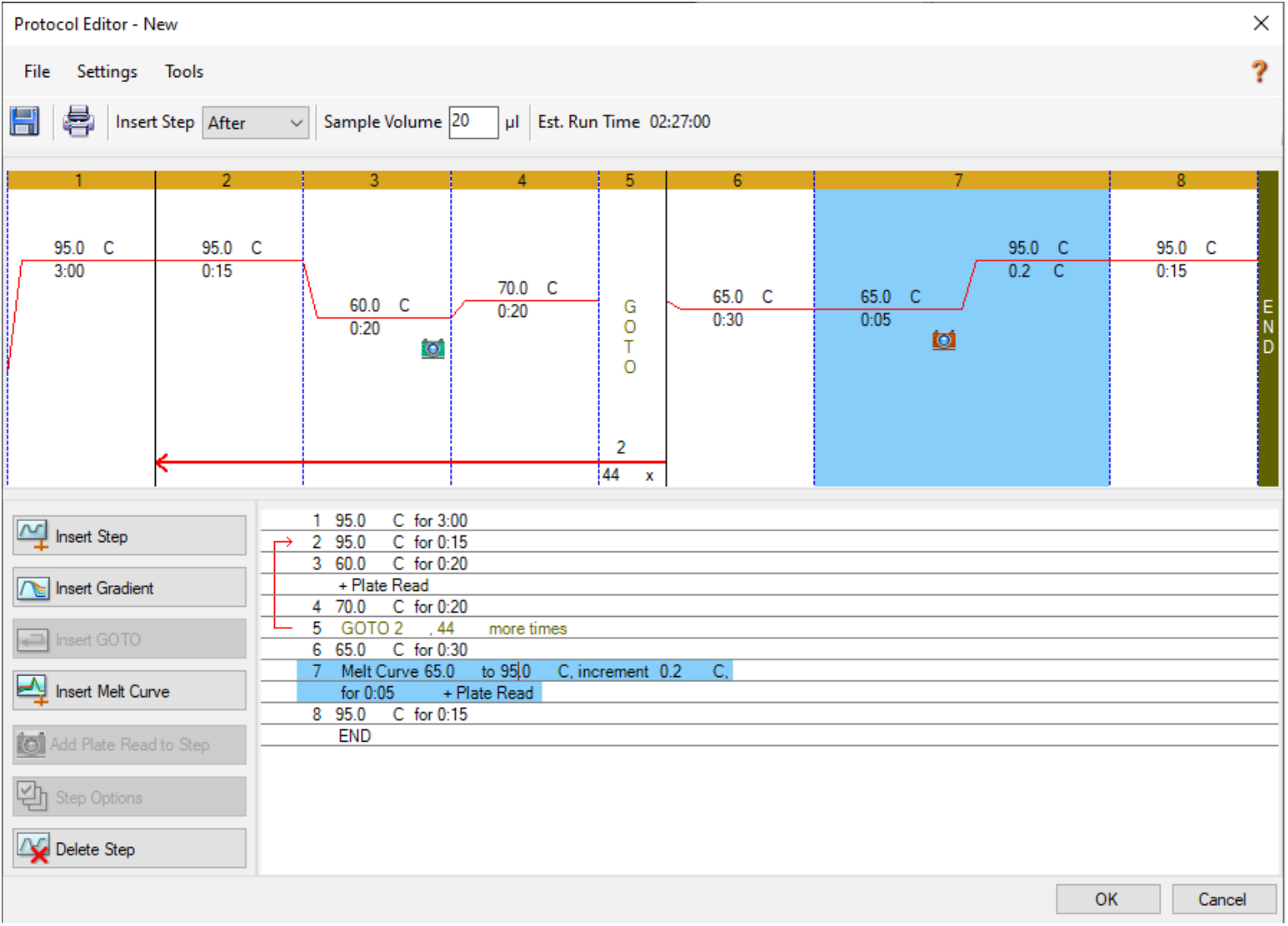

## Analysis

1. Open the .ZPCR file of the run from the real-time PCR machine using CFX Maestro. This will convert the file to a .PCRD, which is the file type accepted by the HRMA software. The Cq value of the negative control can also be checked with Maestro. It should be “N/A” or greater than 38 to signify that there is no contamination.
2. Launch the Precision Melt Analysis software and open the .PCRD file.
  a. File → Open → New → Melt File (.PCRD file)
3. Click on the “View/Edit Plate” button in the toolbar to launch the plate editor view in a new window. This view will be used to exclude all samples except those that are actively being analyzed. This will ensure that the software is only considering the appropriate values that are relevant to the current step in the analysis process, which will prevent other samples from inappropriately influencing the deflection curves.
4. Select all wells **except** the wildtype references and the set of unknown samples without the spike-in. To do this, click and drag while holding the Control key to select the appropriate wells, scroll down on the right panel, and check the “Exclude Wells in Analysis” box (indicated with yellow arrow). Save the changes and exit the plate editor.

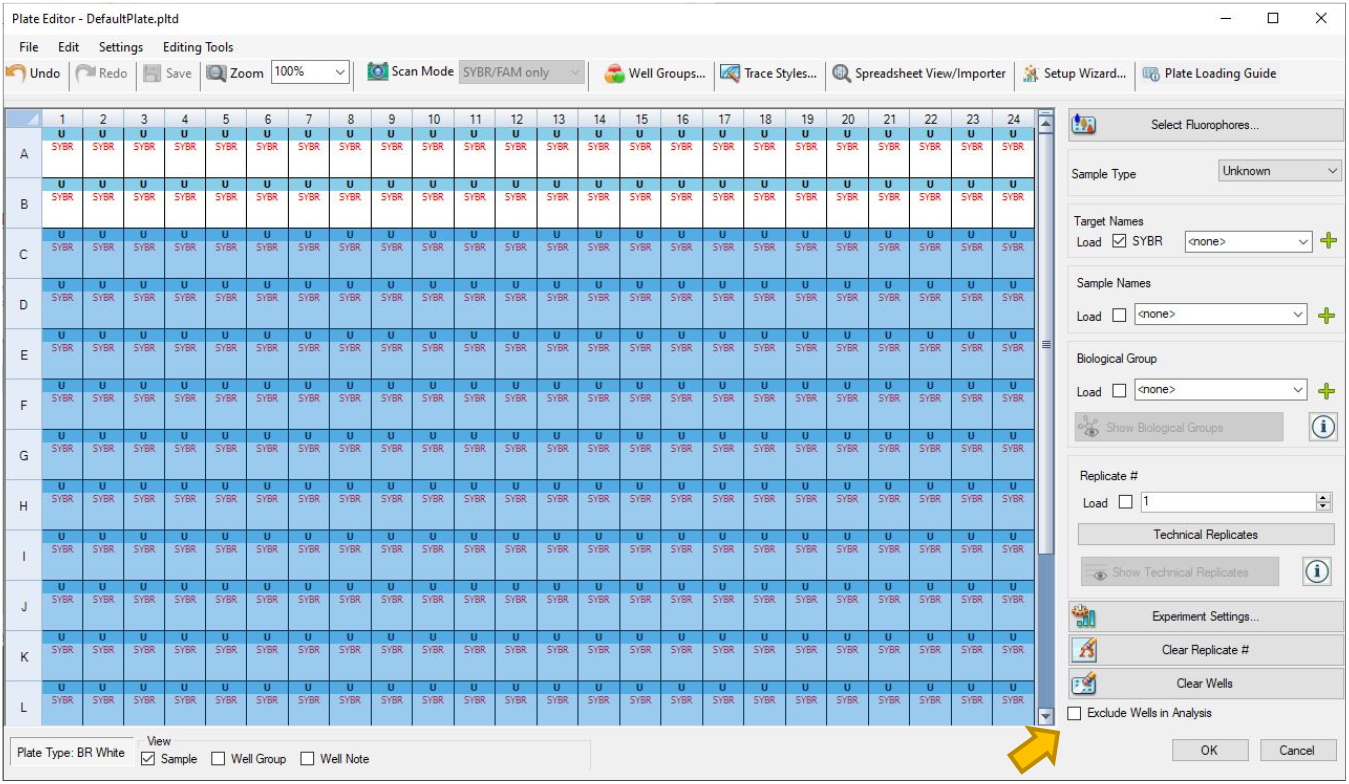
5. The program’s algorithm automatically assigns a reference population. To set the correct reference, gray out all the samples except for the wildtype references in the plate layout view on the bottom left by clicking and dragging to temporarily hide other samples.

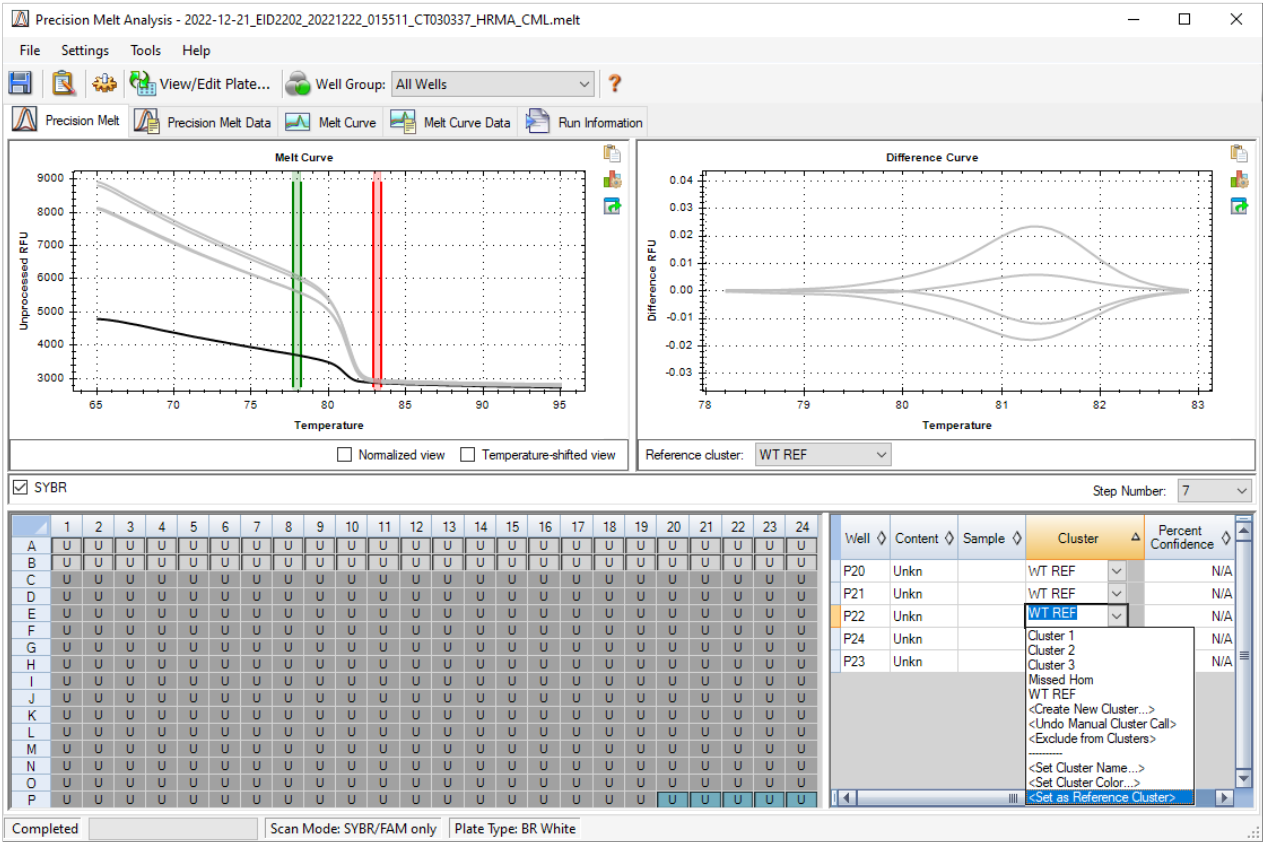
6. Click the dropdown next to the cluster name of one wildtype sample. Select “Create New Cluster” and assign a new name, e.g., *WT ref*. Then, click “Set as Reference Cluster.”
7. Using the same dropdown menu associated with the other wildtype samples and assign each to the same cluster as in Step 7.
  a. Exclude any obvious outliers. Including 5 wildtype reference samples on the plate allows for the exclusion of 1-2 outliers without impacting analysis.
8. Unhide the standard samples from Step 5 (click and drag). Now, the software’s algorithm will automatically assign clusters based upon commonality of melting point deviation from the selected wildtype references.

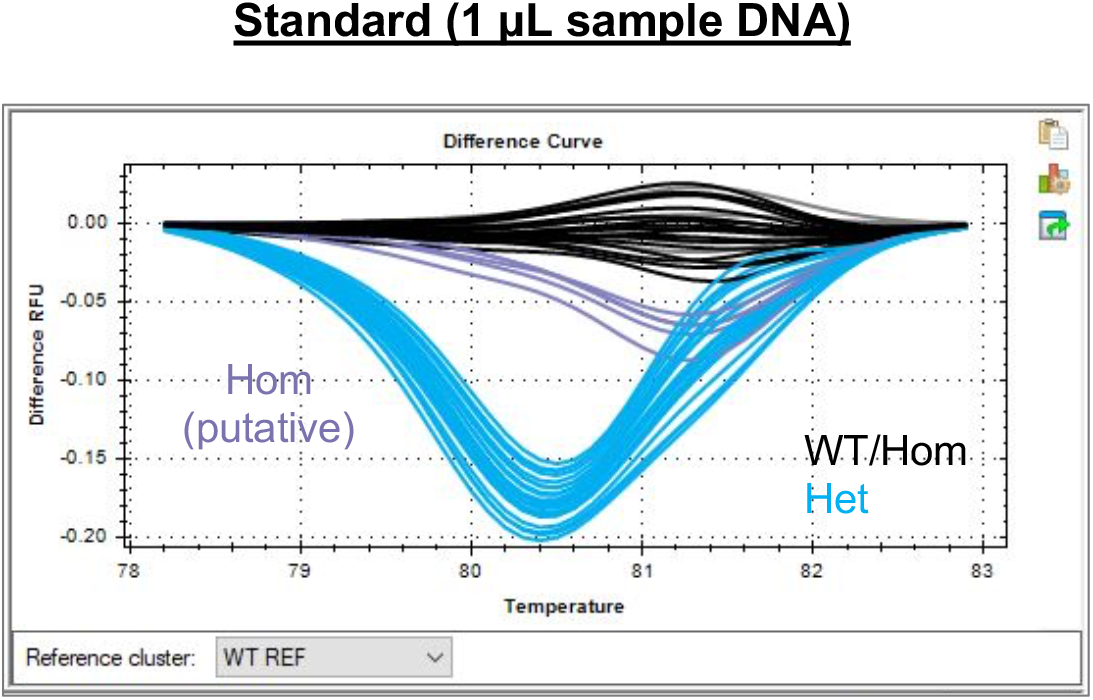
9. All samples with two deflections in the curve are **heterozygotes**. The software will not always group these samples together. Manually assign to the same cluster as needed. The rest of the samples are either wildtypes or homozygotes, which are not reliably distinguishable from one another with only the standard run, although many homozygotes deflect lower than -0.04.
10. (Optional) Create new separate clusters for heterozygotes, wildtypes/homozygotes, and putative homozygotes.
11. To analyze the deflection patterns of the *standard samples*, either:
  a. Hover over the samples in the plate set-up panel to highlight the corresponding deflection curve in the generated graph, OR
  b. Scroll through the samples on the bottom right panel.
12. Using a table similar to the example below, mark samples with a double deflection curve in the standard run, i.e., with an “X” or by coloring in the box. *Optionally, you may choose to call samples with a deflection amplitude lower than -0.04 as putative homozygotes. Regardless, these will be validated with analysis of the corresponding sample with the wildtype spike-in.

13. Return to the plate editor view (View/Edit Plate). Exclude the standard set of samples from analysis and include the set <u>with</u> the wildtype spike-in. Save the changes and exit to the normal view with the four panels displayed.

| Sample ID | Standard | Spike-In | tp53 |
| --- | --- | --- | --- |
| 1 |  |  |  |
| 2 |  |  |  |
| 3 |  |  |  |
| 4 |  |  |  |
| 6 |  |  |  |
| 7 |  |  |  |
| 8 |  |  |  |
| 9 |  |  |  |
| ... |  |  |  |

| KEY |  |
| --- | --- |
| X | Het |
|  | WT/Hom |
| Hom | Hom, Putative |

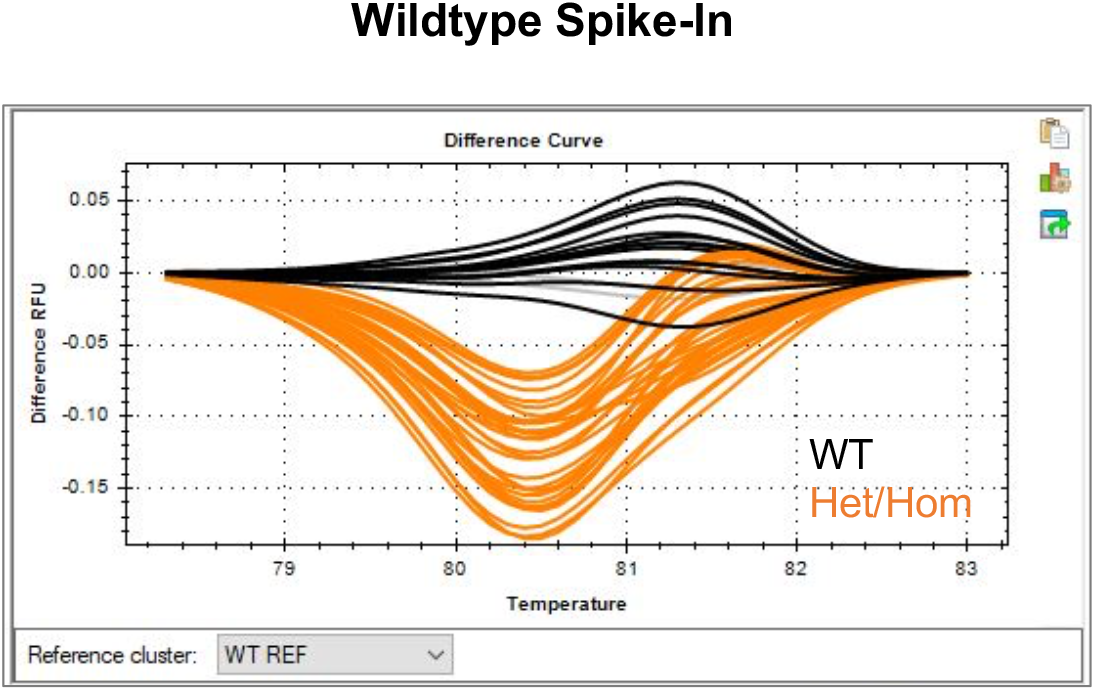
14. This time with the spike-in, the samples with the double deflection are either heterozygotes or homozygotes. Make sure that all samples with this deflection pattern are assigned to the same cluster. All other samples are **wildtypes**.
15. Analyze the deflection patterns of the *spike-in samples* similarly as described above in Steps 11-12, filling in the “Spike-In” column.
16. Compare the results from the analyses of the *standard* and *spike-in* runs using the following key. This parallel comparison strategy allows for *tp53* genotypes to be called with high confidence.

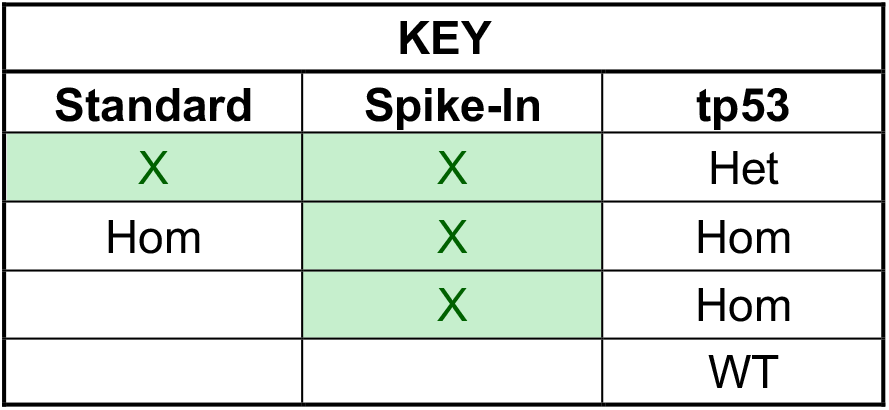

## References

1. Berghmans S, Murphey RD, Wienholds E, et al. tp53 mutant zebrafish develop malignant peripheral nerve sheath tumors. Proc Natl Acad Sci U S A 2005;102(2):407–12, doi:10.1073/pnas.0406252102

2. Kendall GC, Watson S, Xu L, et al. PAX3-FOXO1 transgenic zebrafish models identify HES3 as a mediator of rhabdomyosarcoma tumorigenesis. Elife 2018;7(doi:10.7554/eLife.33800

3. Patton EE, Widlund HR, Kutok JL, et al. BRAF mutations are sufficient to promote nevi formation and cooperate with p53 in the genesis of melanoma. Curr Biol 2005;15(3):249–54, doi:10.1016/j.cub.2005.01.031

4. Olivier M, Hollstein M, Hainaut P. TP53 mutations in human cancers: origins, consequences, and clinical use. Cold Spring Harb Perspect Biol 2010;2(1):a001008, doi:10.1101/cshperspect.a001008

5. White RM, Cech J, Ratanasirintrawoot S, et al. DHODH modulates transcriptional elongation in the neural crest and melanoma. Nature 2011;471(7339):518–22, doi:10.1038/nature09882

6. Zebrafish International Resource Center. tp53zdf1. Eugene, OR; 2008. Available from: http://zebrafish.org/fish/pdf/pcr/zdf1.pdf.

7. Parant JM, George SA, Pryor R, et al. A rapid and efficient method of genotyping zebrafish mutants. Dev Dyn 2009;238(12):3168–74, doi:10.1002/dvdy.22143

8. Talbot JC, Amacher SL. A streamlined CRISPR pipeline to reliably generate zebrafish frameshifting alleles. Zebrafish 2014;11(6):583–5, doi:10.1089/zeb.2014.1047

9. Applied Biosystems. A Guide to High Resolution Melting (HRM) Analysis. Foster City, CA; 2010. Available from: https://tools.thermofisher.com/content/sfs/manuals/cms_070283.pdf.

10. Bertho S, Clapp M, Banisch TU, et al. Zebrafish dazl regulates cystogenesis and germline stem cell specification during the primordial germ cell to germline stem cell transition. Development 2021;148(7), doi:10.1242/dev.187773

